# Large-scale transcriptomic meta-analysis identifies novel therapeutic targets for ulcerative colitis

**DOI:** 10.64898/2026.04.22.719865

**Authors:** Maciej Piernik, Małgorzata Adamiec-Organiściok, Magdalena Skonieczna, Piotr Eder

## Abstract

Ulcerative colitis (UC) is a chronic inflammatory bowel disease for which the vast majority of the approved therapies target the immune system. We aimed to identify consistently dysregulated genes and pathways across independent UC transcriptomic cohorts, distinguishing constitutive from inflammation-dependent changes. We performed a random-effects meta-analysis of 14 microarray datasets from the Gene Expression Omnibus (972 mucosal biopsies, 9 platforms), comparing inflamed UC, uninflamed UC, and inflamed Crohn’s disease (CD) to controls, as well as UC to CD directly. The inflamed UC analysis revealed an upregulated inflammatory transcriptomic profile in UC, providing a rationale for the use of all approved anti-inflammatory therapies. In parallel, the predominant downregulated signal was metabolic, driven by PPARGC1A, PPARGC1B, and ES-RRA, indicating a coordinated, inflammation-dependent collapse. The uninflamed UC analysis revealed a separate set of potentially constitutive vulnerabilities — fully suppressed glucuronidation, upregulated translation, complement priming, and altered iron export — that are not downstream of the energy collapse. The metabolic deficit was more severe in UC than in CD, while immune pathways were shared. These findings suggest a two-layer model of UC pathology: a constitutive impairment of metabolic pathways that is further exacerbated by inflammation. The inflammation analysis reveals new targets for immune suppression while the constitutive analysis identifies targets for proactive intervention between flares directed at the metabolic deficiency.

## Introduction

Ulcerative colitis (UC) is a chronic inflammatory bowel disease (IBD) characterized by mucosal inflammation of the colon, typically beginning in the rectum and extending continuously (1). The disease follows a relapsing-remitting course and its clinical hallmarks are rectal bleeding and bloody diarrhea (1). The global incidence of UC has been steadily increasing, particularly in newly industrialized countries, suggesting that environmental and lifestyle factors play a significant role in disease development (2).

Currently, there is no cure for UC. Available therapies are largely assigned empirically as there are no reliable biomarkers determining which patients will respond to specific treatments (3, 4). Nearly all approved therapies function primarily as immunosuppressants, targeting various components of the inflammatory system rather than the underlying disease cause. Despite significant advancements in treatment options for UC in the recent years, with the advent of an increasing number of targeted therapies that mute the immune response (5–11), treatment outcomes remain unsatisfactory. Primary non-response occurs in 20–40% of patients with UC receiving biologic therapy (12–14) while the secondary loss of response among initial responders develops at an annual rate of 10–23% during anti-TNF maintenance therapy, with the highest risk in the first year (12, 15). When medical therapy fails or in cases of severe complications such as toxic megacolon or cancer, total colectomy remains the only definitive treatment, performed in approximately 10% of UC patients within 10 years of diagnosis (16, 17).

The fundamental reason for the lack of curative therapy is that, despite extensive research, the exact etiology of UC remains unknown (18–20). Genome-wide association studies have identified over 240 risk loci related to IBD (21), including 120 UC-specific (22), yet these explain only a fraction of disease heritability (21). The continued rise and geographic expansion of UC (2) occurring on timescales too short to explain by genetic change (23) highlight the importance of environmental and metabolic factors that remain poorly characterized (24, 25).

Gene expression studies of intestinal mucosal biopsies have provided valuable insights into the molecular mechanisms underlying UC (26–28). Transcriptomic profiling enables characterization of the disease at the site of inflammation, revealing alterations in immune, inflammatory, and metabolic pathways (26, 29, 30). Over the past two decades, thousands of transcriptomic studies have deposited intestinal and other gene expression datasets in public repositories such as GEO (31), creating an unprecedented resource for large-scale analysis. This accumulation of datasets enables meta-analytic approaches that can identify consistently dysregulated genes across independent cohorts, distinguishing real disease signatures from study-specific artifacts and biases (32). Such cross-validation across multiple datasets offers a unique opportunity to identify the strongest biological signals that may point to core disease mechanisms.

The aim of this study was to conduct a comprehensive meta-analysis of all available microarray gene expression data from mucosal biopsies of UC patients and controls in order to identify consistently dysregulated genes and pathways that may contribute to disease etiology and pathogenesis. By integrating data across 14 datasets (972 samples) from the Gene Expression Omnibus, our goal was to identify molecular signatures that bypass the limitations of any individual study.

## Materials and Methods

The overall study design and analytical workflow presented in this section are summarized in Figure 1.

**Fig. 1.**
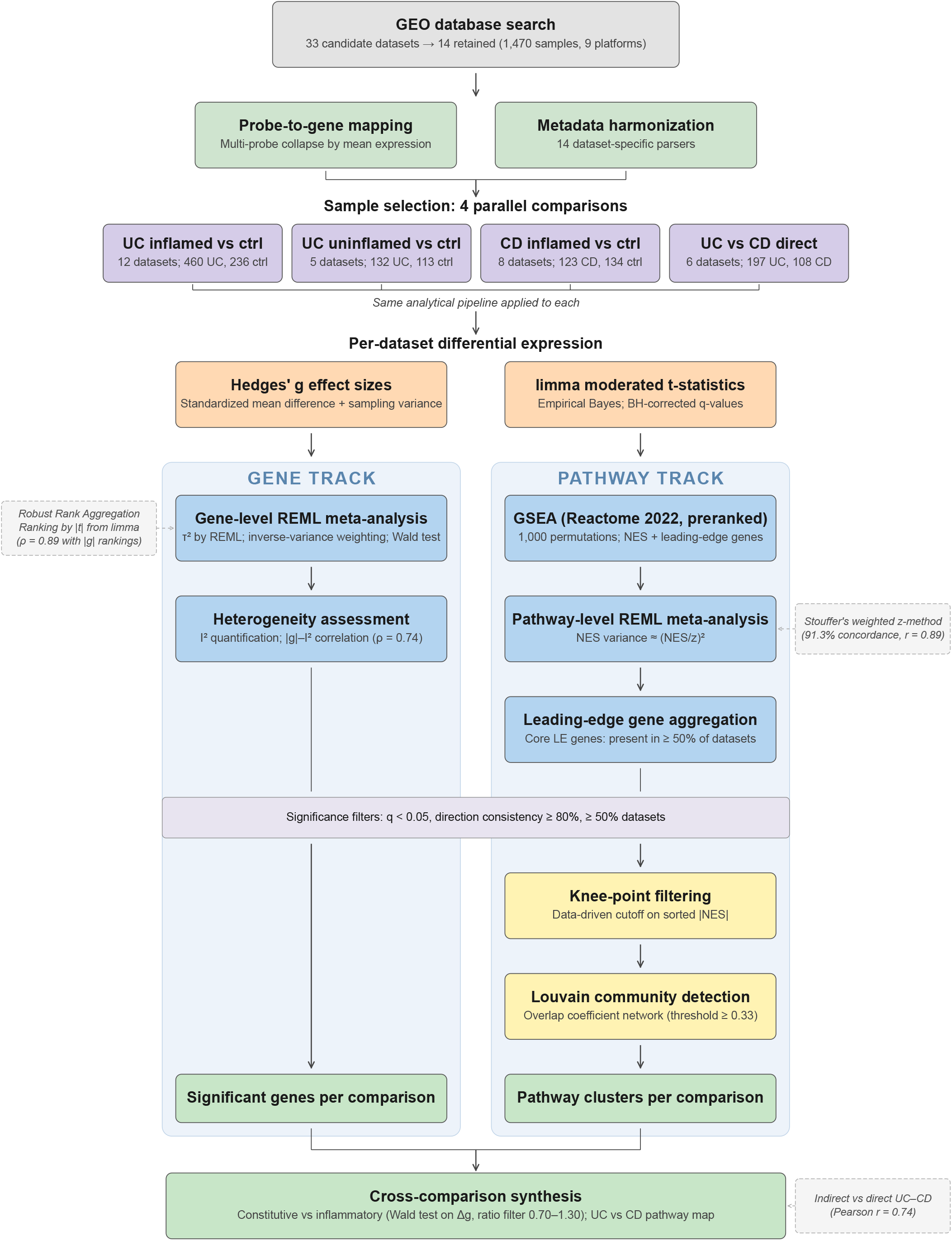
Overview of the analytical pipeline. Dashed boxes indicate analyses performed to cross-validate the primary results.

### Data

We searched the NCBI Gene Expression Omnibus (GEO) (31) for microarray gene expression datasets derived from intestinal mucosal biopsies of patients with IBD. The search was performed using the terms “ulcerative colitis” and “inflammatory bowel disease”, restricted to *Homo sapiens* expression profiling by array. We identified 33 candidate datasets encompassing a total of 2,456 samples.

Datasets were included if they (i) contained gene expression data from colonic or rectal mucosal biopsies, (ii) were generated on a microarray platform for which probe-to-gene annotation was available, and (iii) contained samples that could be unambiguously classified by disease status based on the metadata. Datasets were excluded if they represented subsets, combinations, or duplicates of other studies.

After applying these criteria, 14 independent datasets containing a total of 1,470 samples were retained, spanning 9 microarray platforms (Table 1). Sample-level inclusion criteria (described below) further reduced this to 972 samples across the four analyses. The included datasets comprised samples from patients with UC, Crohn’s disease (CD), and controls, with a subset additionally including treatment and response information.

**Table 1.** Dataset characteristics; *n*_*UC*_ -inflamed UC samples, *n*_*uninfl*_ -un-inflamed UC samples, *n*_*CD*_ -inflamed CD samples, *n*_*Ctrl*_ -uninflamed control samples.

| Dataset | Platform | $n_{UC}$ | $n_{uninfl}$ | $n_{CD}$ | $n_{Ctrl}$ |
| --- | --- | --- | --- | --- | --- |
| GSE11223 | GPL1708 | 62 | 61 | - | 63 |
| GSE16879 | GPL570 | 24 | - | 37 | 12 |
| GSE179285 | GPL6480 | 23 | 32 | 14 | 23 |
| GSE22619 | GPL570 | 10 | - | - | 10 |
| GSE36807 | GPL570 | 15 | - | 13 | 7 |
| GSE48634 | GPL10558 | 56 | - | 29 | 56 |
| GSE53306 | GPL14951 | 16 | 12 | - | 12 |
| GSE6731 | GPL8300 | 5 | 4 | 7 | 4 |
| GSE73661 | GPL6244 | 67 | - | - | 12 |
| GSE75214 | GPL6244 | 74 | 23 | 8 | 11 |
| GSE92415 | GPL13158 | 87 | - | - | 21 |
| GSE9452 | GPL570 | 21 | - | - | 5 |
| GSE52746 | GPL17996 | - | - | 10 | 17 |
| GSE72780 | GPL570 | - | - | 5 | 4 |
| **Total** |  | **460** | **132** | **123** | **257** |

### Data processing

A custom data processing pipeline was implemented in Python to integrate the heterogeneous microarray datasets. Raw expression data and sample meta-data were downloaded programmatically from GEO using the GEOparse library (33). Processing consisted of two main steps: probe-to-gene mapping and metadata standardization.

### Probe-to-gene mapping

For each microarray platform, probe identifiers were mapped to NCBI Entrez Gene IDs using the platform-specific annotation tables deposited in GEO. The mapping procedure was adapted to the annotation format of each platform (details in the publicly available processing code). Probes that could not be mapped to any Entrez Gene ID were excluded. When multiple probes mapped to the same gene within a dataset, expression values were collapsed by mean expression, yielding a gene-level expression matrix for each dataset.

### Metadata standardization

Sample metadata in GEO are stored in a key-value text format with no enforced schema, resulting in substantial heterogeneity across datasets. To enable cross-dataset comparison, a dataset-specific metadata processor was implemented for each of the 14 included studies, extracting and unifying disease status (UC, CD, or control), treatment, treatment response, tissue location, inflammation status, and endoscopic Mayo score where available. For datasets from interventional studies, treatment response was coded as responder or non-responder based on the criteria defined in the original publications.

No additional cross-study normalization was applied beyond what was performed in the original study — each dataset was used as provided after probe-to-gene mapping. Because effect sizes were estimated separately within each dataset as standardized mean differences (Hedges’ *g*), study-specific differences in measurement scale or baseline expression were largely mitigated. Remaining between-study heterogeneity was then modeled, rather than removed, using a random-effects meta-analysis. This design was chosen deliberately to avoid the assumptions inherent in cross-platform normalization methods, which can introduce their own artefacts when applied to heterogeneous microarray data. All processing code is publicly available at https://github.com/MaciejPiernik/ibd.

### Ethical statement

This study used only publicly available, de-identified gene expression data from the Gene Expression Omnibus. No new human samples were collected. All original studies obtained appropriate ethical approvals as described in their respective publications.

### Sample selection

Four comparative analyses were performed, each applying the same analytical pipeline (per-dataset differential expression, gene-level and pathway-level meta-analysis, knee-point filtering, and community detection — all detailed later in this section). In all cases, datasets with fewer than 6 total samples or fewer than 3 samples in either group after selection were excluded. To ensure expression values were on comparable scales across platforms, the median expression level of each dataset was checked at loading time. Datasets with a median exceeding 100 (indicating non-log-transformed intensities) were transformed by applying log_2_(*x* + 1).

### UC inflamed vs controls (primary analysis)

We compared inflamed colonic mucosal biopsies from UC patients with uninflamed colonic biopsies from controls (non-IBD patients).

Samples were included if they were (i) from colonic or rectal tissue (ileal biopsies were excluded), (ii) obtained prior to or at baseline of any intervention (treatment-naive or week 0 / pre-treatment in interventional studies), and (iii) classifiable as either UC with active inflammation or control without inflammation based on the harmonized metadata. Where inflammation status was not explicitly annotated, UC samples were assumed inflamed and control samples uninflamed.

### UC uninflamed vs controls

To distinguish constitutive disease-associated changes from those driven by active inflammation, we compared uninflamed UC biopsies with uninflamed controls. The same inclusion criteria as the primary analysis were applied, except that UC samples were required to be annotated as uninflamed.

### CD inflamed vs controls

To compare UC with CD, we identified inflamed colonic CD biopsies from datasets that also contained controls. The same tissue, treatment, and inflammation criteria were applied.

### Direct UC vs CD comparison

As an additional validation, we directly compared inflamed UC and inflamed CD biopsies within the datasets that contained both disease groups. This design entirely omits controls, so that UC–CD differences are estimated within each study without relying on each disease’s deviation from a common control group.

### Per-dataset differential expression

For each dataset, two parallel streams of gene-level statistics were computed.

### Effect sizes for meta-analysis

For each gene, group means, variances, and sample sizes were computed for the UC and control groups. The standardized mean difference was calculated as Hedges’ *g*, a bias-corrected version of Cohen’s *d*:

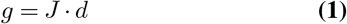

where

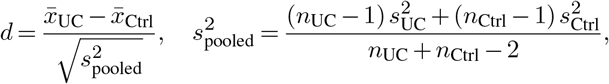

and *J* = 1−3*/*(4(*n*_UC_ + *n*_Ctrl_) − 9) is the small-sample correction factor (34). The approximate sampling variance of Hedges’ *g* was estimated as *v* = *n/*(*n*_UC_·*n*_Ctrl_) + *g*^2^*/*(2*n*), where *n* = *n*_UC_ + *n*_Ctrl_ (35). These per-gene variance estimates served as the within-study variances *v*_*i*_ in the subsequent meta-analysis.

### Ranking statistics for gene set enrichment

Moderated *t*-statistics were obtained using the limma empirical Bayes framework (36, 37) via the R limma package through rpy2. For each gene, limma fits a linear model comparing UC and controls. It then stabilizes the gene-specific noise estimates by borrowing information across all genes, which improves statistical reliability, especially in small datasets. The resulting moderated *t*-statistics were used to rank genes for gene set enrichment analysis, and the associated *p*-values were corrected for multiple testing using the Benjamini-Hochberg procedure (38) to obtain per-dataset *q*-values.

### Gene-level meta-analysis

Gene-level effect sizes (Hedges’ *g*) were combined across datasets using a random-effects model. For each gene measured in at least two datasets, the observed effect in dataset *i* is modelled as *y*_*i*_ ∼𝒩 (*µ, τ*^2^ + *v*_*i*_), where *µ* is the overall true effect, *v*_*i*_ the within-study sampling variance, and *τ*^2^ the between-study variance. *τ*^2^ was estimated by restricted maximum likelihood (REML) (39) using the PyMARE library (40), and the combined effect computed as 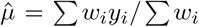, where 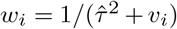, with standard error SE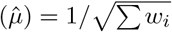. Statistical significance was assessed by Wald test 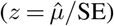with Benjamini–Hochberg correction. Heterogeneity was quantified by *I* ^2^ (41).

For each gene, the direction of effect was classified as upregulated or downregulated based on the sign of the majority of per-dataset effect sizes, and a direction consistency ratio was computed as the fraction of datasets agreeing on the majority direction. A gene was considered significant if it met all of the following criteria: *q* < 0.05, direction consistency ≥0.8, and presence in at least 50% of datasets in the given analysis (i.e. ≥ 6 of 12 for inflamed UC, ≥ 3 of 5 for uninflamed UC, ≥ 4 of 8 for CD, and ≥ 3 of 6 for direct UC vs CD).

### Robust Rank Aggregation

As a complementary non-parametric approach, gene rankings were aggregated across datasets using Robust Rank Aggregation (RRA) (42). Within each dataset, genes were ranked by the absolute value of the moderated *t*-statistic and ranks were normalized to [0, 1] by dividing by the total number of genes tested. Under the null hypothesis of no differential expression, each gene’s normalized rank follows a uniform distribution on [0, 1]. For a gene observed across *k* datasets with sorted normalized ranks *u*_(1)_ *≤ · · · ≤ u*_(*k*)_, the RRA *ρ* statistic was computed as:

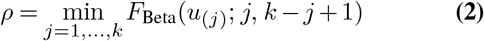

where *F*_Beta_ denotes the cumulative distribution function of the Beta distribution. A small *ρ* indicates that the gene ranks consistently higher across datasets than expected by chance. The *p*-value was approximated as *p* ≈ min(1, *ρ*·*k*) (42), and *q*-values were obtained using the Benjamini–Hochberg procedure.

### Gene set enrichment analysis

Gene set enrichment analysis (GSEA) (43) was performed independently in each dataset using the preranked method implemented in GSEApy (44). Genes were ranked by their moderated *t*-statistic (most upregulated first) and tested against the Reactome 2022 pathway database (45). Enrichment was quantified by the standard running-sum statistic and normalized to yield the normalized enrichment score (NES) (43). Nominal *p*-values were estimated from 1,000 gene-set permutations, and false discovery rates were computed across all tested pathways. Pathways with fewer than 5 or more than 1,000 genes were excluded. For each pathway, the leading-edge gene set — the genes driving the enrichment signal — was recorded in each dataset.

### Pathway-level meta-analysis

Pathway NES values were combined across datasets using the same REML random-effects framework as for gene-level analysis. Because direct sampling variance is not provided for NES, it was approximated from the *p*-value assuming normality: 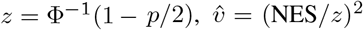 (35). This approximation was validated by a sensitivity analysis using Stouffer’s weighted *z*-method (see Results). The same filtering criteria were applied: *q* < 0.05, direction consistency ≥ 0.8, presence in ≥ 50% of datasets.

### Leading-edge gene aggregation

For each significant pathway, leading-edge genes were aggregated across datasets. A gene was designated as a “core leading-edge” gene if it appeared in the leading edge in at least half of the datasets with available data.

### Pathway selection and clustering

#### Knee-point filtering

To focus the downstream interpretation on the most strongly supported signals, an additional prioritization step was applied. Instead of analyzing an arbitrary fixed number of top-ranked pathways, a data-driven cutoff was defined from the knee point of the sorted |*NES*| distribution. Pathways were sorted by descending |*NES*|, and the knee point was detected using the knee locator algorithm (46) as implemented in the kneefinder Python package, which identifies the point of maximum deviation from the straight line connecting the best and worst score in the sorted |*NES* |curve. Pathways beyond this point were not carried forward for downstream interpretation.

#### Community detection

To reduce redundancy among prioritized pathways and identify broader functional motifs, a network was constructed in which nodes represent significant pathways and edges represent gene overlap. For each pair of pathways, the overlap coefficient was computed as |*A*∩*B*| */* min(|*A, B*|), where *A* and *B* are the core leadingedge gene sets of the two pathways. Edges with overlap coefficient below 0.33 were discarded. Community structure was detected using the Louvain algorithm (47), which partitions the network into groups of pathways sharing more genes internally than with other groups. Each community was characterised by its constituent pathways and mean NES.

## Results

### Dataset overview

12 gene expression datasets with inflamed colonic mucosal biopsies from UC patients and controls (non-IBD patients) met the inclusion criteria for the main analysis (Table 1). The combined cohort consisted of 460 UC and 236 control samples across 8 microarray platforms. Per-dataset sample sizes ranged from 9 (GSE6731) to 125 (GSE11223). The number of differentially expressed genes (DEGs; limma, *q* < 0.05) varied significantly between datasets, from 2 in GSE22619 to 10,872 in GSE92415, partly reflecting differences in sample sizes. 3 additional analyses were performed in parallel: uninflamed UC versus controls (5 datasets; 132 UC, 113 controls), inflamed CD versus controls (8 datasets; 123 CD, 134 controls), and a direct UC versus CD comparison within 6 datasets containing both disease groups (197 UC, 108 CD). A summary of all 4 analyses is provided in Table 2.

**Table 2.**
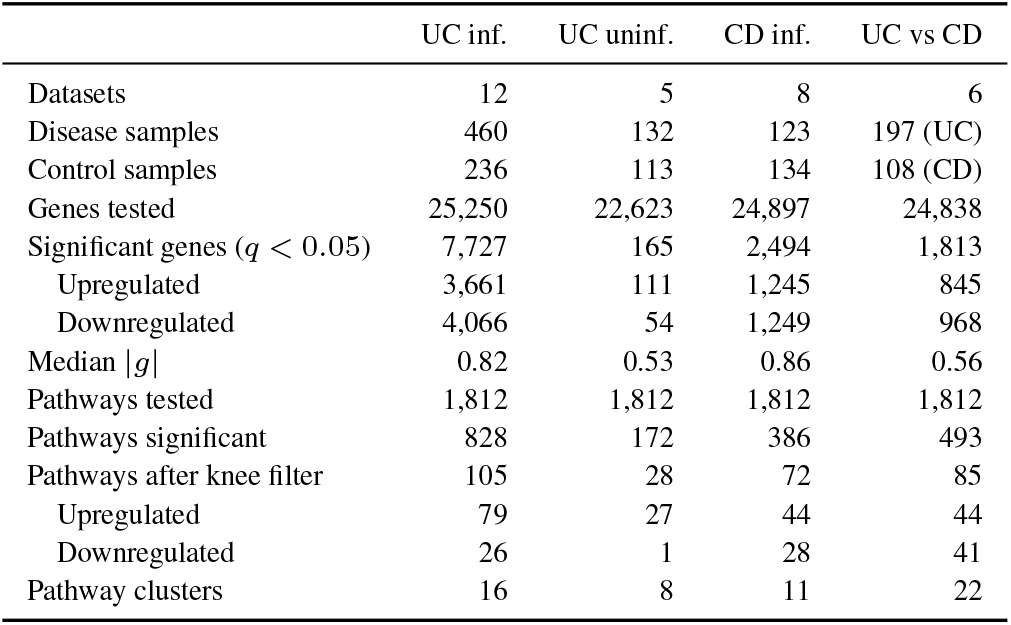
Summary of the four parallel meta-analyses. For the three disease vs control comparisons, upregulated/downregulated refers to the direction in disease relative to controls. For the direct UC vs CD comparison, upregulated means higher in UC and downregulated means lower in UC relative to CD. Significant genes: random-effects meta-analysis, *q* < 0.05, direction consistency *≥* 80%. Pathway filtering: knee-point on ranked |NES|; clusters: Louvain community detection on shared leading-edge genes.

### Gene-level meta-analysis of inflamed UC

Random-effects meta-analysis identified 7,727 significantly dysregulated genes out of 25,250 tested (*q* < 0.05, direction consistency ≥80%, presence in ≥50% datasets), of which 3,661 were upregulated and 4,066 downregulated. The median absolute effect size among significant genes was |*g*| = 0.82, with a median heterogeneity of *I* ^2^ = 0.76 (Supplementary Figure S1A). This level of heterogeneity is expected in a cross-platform meta-analysis integrating 8 microarray technologies and reflects variation in the magnitude of dysregulation across cohorts (driven by differences in sample sizes, disease severity, and platform sensitivity) rather than inconsistency in direction, which was controlled by the ≥ 80% direction consistency filter. Heterogeneity was strongly associated with effect size (Spearman *ρ* = 0.74 between |*g*| and *I* ^2^; Supplementary Figure S1B): the top 100 genes ranked by |*g*| had a median *I* ^2^ = 0.93, while the top 100 ranked by *q*-value had a median *I* ^2^ = 0.36. These two rankings capture complementary properties: ranking by effect size identifies genes with the largest average dysregulation (e.g. LPCAT1, *g* = +3.14, *I* ^2^ = 0.96), while ranking by statistical significance favours genes with highly consistent, reproducible effects across all cohorts (e.g. ANO5, *g* = −1.62, *I*^2^ = 0). Forest plots for representative genes illustrating this range of heterogeneity are provided in Supplementary Figure S2. Rankings were validated by Robust Rank Aggregation (Spearman *ρ* = 0.89 between |*g*| and RRA *ρ* rankings). Figure 2 visualizes the gene-level results of this analysis while full gene-level results for all analyses are provided in Supplementary Tables S1–S4.

**Fig. 2.**
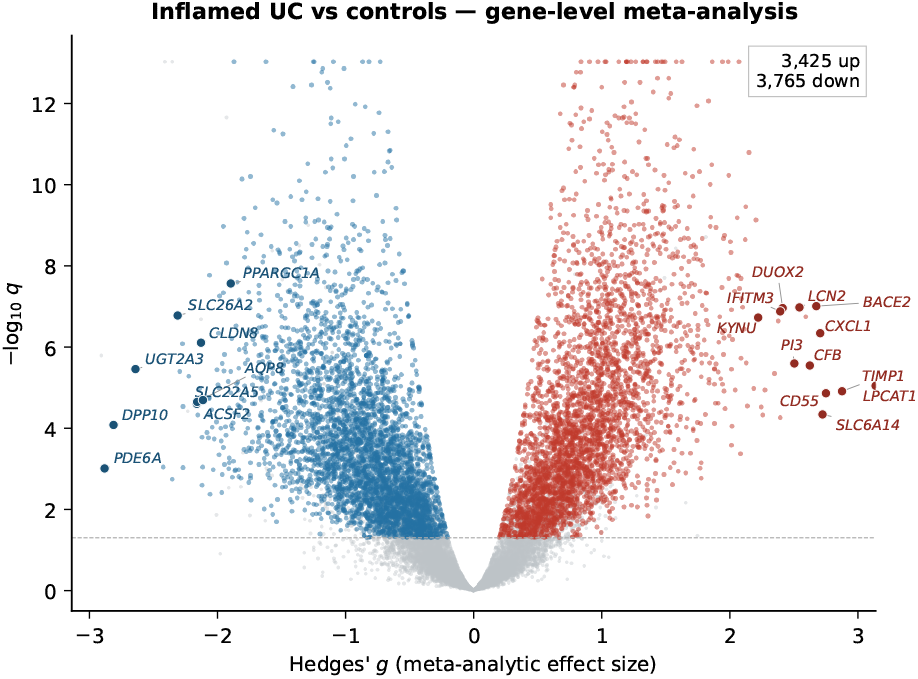
Volcano plot for inflamed UC versus controls. Each point represents a protein-coding gene. Axes show meta-analytic Hedges’ _*g*_(horizontal) and *−* log_10_ *q* (vertical); the dashed line marks *q* = 0.05. Color indicates significance (*q* < 0.05, direction ratio *≥* 0.8, *≥* 50% dataset coverage), with red denoting up-regulation, blue denoting downregulation, and grey denoting non-significant genes. Gene names are shown for selected top extremes and key discussion genes.

The most strongly upregulated genes (Table 3) included LPCAT1 (*g* = +3.14), TIMP1 (*g* = +2.88), CD55 (*g* = +2.75),and SLC6A14 (*g* = +2.72). Unsurprisingly, most of the top upregulated genes mapped to immune-/inflammation-related pathways: LPCAT1 and S100A11 (neutrophil degranulation), TIMP1, CXCL1, LCN2, IFITM3, UBE2L6 (cytokine signaling), CD55, CFB, C2 (complement system), DUOX2, DUOXA2 (oxidative stress). Very few of the upregulated genes were not assigned to any pathway, and out of those only BACE2 and RBPMS are not directly immune-related. Among downregulated genes (Table 3), the largest effects were observed for PDE6A (*g* = −2.88), DPP10 (*g* = −2.81), and UGT2A3 (*g* = −2.64). More broadly, several of the most strongly downregulated genes pointed to disrupted metabolism and epithelial homeostasis, including ACSF2 (acyl-CoA synthetase, fatty acid metabolism), SLC22A5 (carnitine transporter, fatty acid metabolism), UGT2A3 (glucuronidation), and AQP8 (aquaporin 8, colonocyte water transport). Notably, many of the most strongly downregulated genes were not assigned to any of the top enriched pathways, suggesting that some of the most extreme metabolic changes occur in genes outside canonical pathways. Reporting in Table 3 was restricted to protein-coding genes; non-coding RNAs and uncharacterised loci were left in the meta-analysis but are not discussed individually.

**Table 3.**
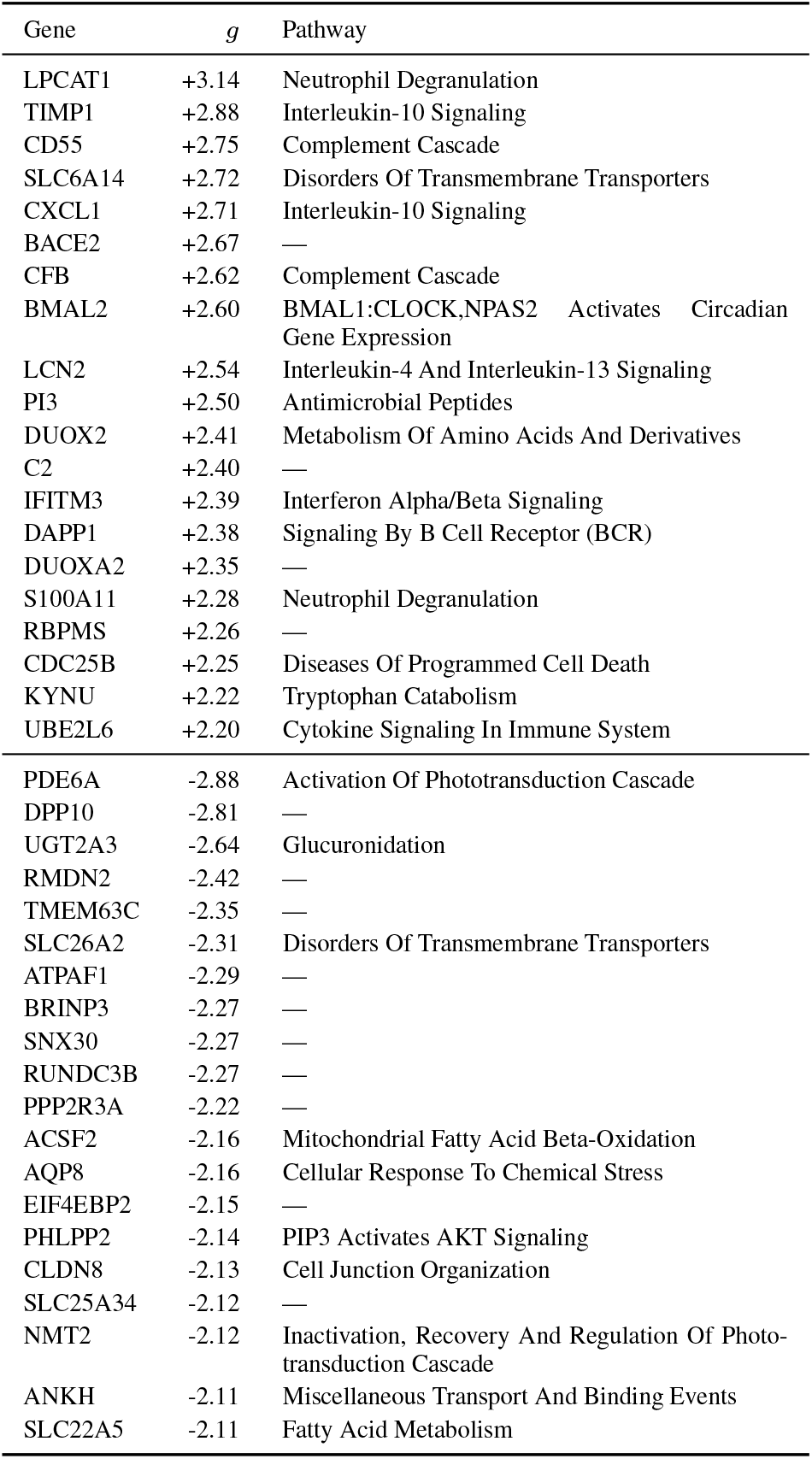
Top 20 up/downregulated genes by Hedges’ *g* from random-effects meta-analysis of inflamed UC vs control samples. Pathway indicates the highest-scoring enriched Reactome pathway containing the gene in its leading edge; “—” denotes genes not present in any enriched pathway.

### Pathway enrichment and clustering

Gene set enrichment analysis of Reactome pathways, followed by pathway-level random-effects meta-analysis, identified 828 significantly enriched pathways out of 1,812 tested. As a sensitivity check, an independent Stouffer weighted *z*-method analysis (combining per-dataset *p*-values weighted by 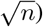confirmed 91.3% of REML-significant pathways at *q* < 0.05, with 100% direction concordance and Pearson *r* = 0.89 between NES and Stouffer *z* (Supplementary Figure S3), supporting the appropriateness of the NES variance approximation. Knee-point filtering reduced this set to 105 pathways (79 upregulated, 26 downregulated), which were grouped into 16 communities by Louvain clustering on shared leading-edge genes (Table 4; full pathway results for all analyses are provided in Supplementary Tables S5–S12).

**Table 4.**
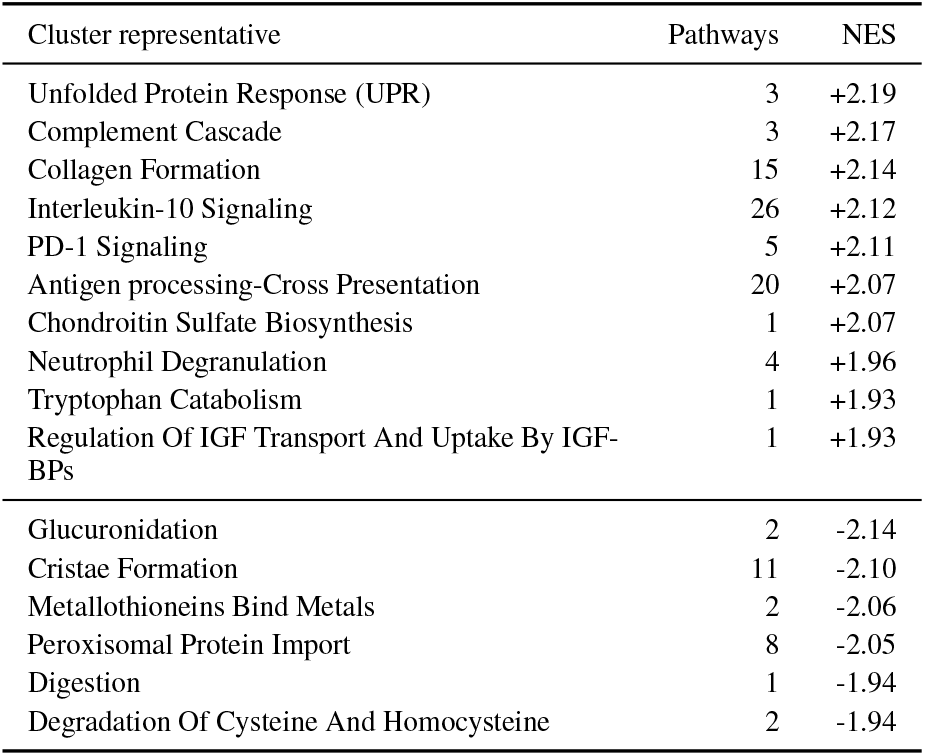
Pathway clusters in inflamed UC. Each row is a community of co-enriched Reactome pathways identified by Louvain clustering on shared leading-edge genes. Pathways: number of member pathways; NES: mean normalized enrichment score (positive = upregulated in UC).

Three broad modules dominated the upregulated pathways: cytokine signaling (represented by Interleukin-10 Signaling, 26 pathways, mean NES = +2.12), antigen processing and adaptive immune signaling (represented by Antigen processing-Cross Presentation, 20 pathways, mean NES = +2.07), and extracellular matrix organisation (represented by Collagen Formation, 15 pathways, mean NES = +2.14). Smaller upregulated clusters captured PD-1/T-cell co-stimulation (represented by PD1-Signaling, 5 pathways, mean NES = +2.11), Complement Cascade (3 pathways, mean NES = +2.17), Unfolded Protein Response (3 pathways, mean NES = +2.19), and Neutrophil Degranulation (4 pathways, mean NES = +1.96), alongside several single-pathway signals.

Among downregulated pathways, the dominating clusters were mitochondrial energy metabolism (represented by Cristae Formation, 11 pathways, mean NES = *−*2.10) and fatty acid/peroxisomal metabolism (represented by Peroxisomal Protein Import, 8 pathways, mean NES = −2.05). Smaller downregulated clusters included glucuronidation (2 pathways, mean NES = *−*2.14), metallothionein-mediated metal binding (2 pathways, mean NES = *−*2.06), sulfur amino acid metabolism (2 pathways, mean NES = *−*1.94), and digestion (1 pathway, NES = *−*1.94).

### Constitutive and inflammation-dependent changes

To distinguish gene expression changes intrinsic to the UC mucosa from those driven by active inflammation, we repeated the meta-analysis using uninflamed UC biopsies versus controls. 5 datasets (GSE11223, GSE179285, GSE53306, GSE6731, GSE75214) contained uninflamed UC samples, yielding 132 UC and 113 control samples (see Table 1). This analysis identified 165 significantly dysregulated genes (*q* < 0.05, present in ≥50% datasets), of which 117 were also significant in the inflamed analysis, all 117 showing the same direction of change. The overall correlation of effect sizes across all 22,623 shared genes was moderate (Pearson *r* = 0.54), reflecting the attenuation of most inflammatory signals in uninflamed mucosa.

To formally classify genes by their inflammation dependence, a Wald test was applied to the difference in meta-analytic effect sizes between the inflamed and uninflamed analyses for each gene present in both 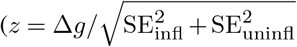, Benjamini–Hochberg corrected). Among the 117 genes significant in both analyses, 96 had no significant difference between conditions (*q*_Δ_≥0.05) and 21 were classified as inflammation-amplified (*q*_Δ_ < 0.05; Supplementary Tables S13–S14). To further select constitutive genes from those that were simply not significantly different, we applied an additional ratio filter (0.70 ≤*g*_uninfl_*/g*_infl_≤1.30), yielding 27 constitutive genes (17 upregulated, 10 downregulated), illustrated in Figure 3.

**Fig. 3.**
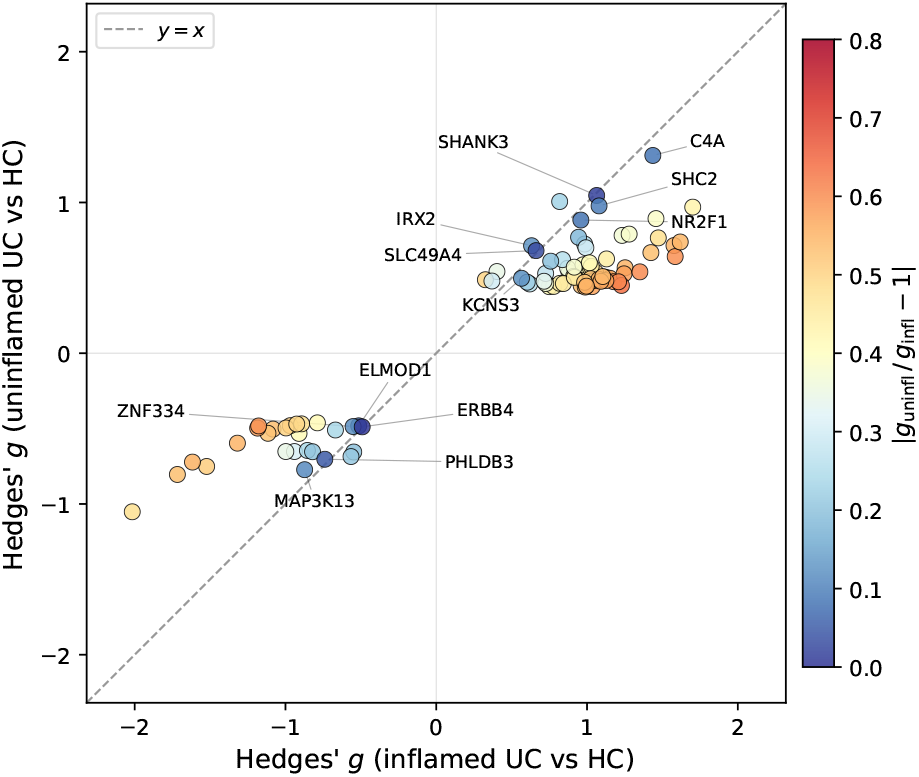
Effect-size comparison for constitutive genes. Each point represents a gene significant in both the inflamed vs ctrl and uninflamed vs ctrl meta-analyses with no significant difference between conditions (Wald *q*_Δ_ *≥* 0.05, *n* = 96). Axes show meta-analytic Hedges’ *g* in the inflamed (horizontal) and uninflamed (vertical) comparisons; the dashed line marks *y* = *x* (identical effect sizes). Color indicates |*g*_uninfl_*/g*_infl_ *−* 1|, with dark blue denoting near-equal effects and warm tones denoting attenuation in uninflamed tissue. Gene names are shown for the subset with ratio 0.85–1.15.

Among the 17 constitutively upregulated genes, C4A showed the largest uninflamed effect (*g*_uninfl_ = +1.31, ratio = 0.91), followed by the scaffolding protein SHANK3 (*g*_uninfl_ = +1.05, ratio = 0.98), the embigin cell-surface glycoprotein EMB (*g*_uninfl_ = +1.01, ratio = 1.23), and the signaling adaptor SHC2 (*g*_uninfl_ = +0.98, ratio = 0.91). Additional genes included the transcription factor NR2F1 (*g*_uninfl_ = +0.88, ratio = 0.92), the extracellular matrix glycoprotein SPARCL1 (*g*_uninfl_ = +0.70, ratio = 0.70), and the iron exporter SLC40A1 (ferroportin; *g*_uninfl_ = +0.62, ratio = 0.74). The 10 constitutively downregulated genes were led by MAP3K13 (*g*_uninfl_ = −0.77, ratio = 0.89) and PHLDB3 (*g*_uninfl_ = −0.70, ratio = 0.95). WDR1 (*g*_uninfl_ = −0.69, ratio = 1.21) and DLGAP1 (*g*_uninfl_ = − 0.66, ratio = 1.20) showed slightly larger uninflamed than inflamed effects. PANK4 (pantothenate kinase 4; *g*_uninfl_ = −0.65, ratio = 0.80), which appeared in the coenzyme A biosynthesis leading edge, was the only core metabolic pathway gene to meet the constitutive ratio criterion. GLIPR2 (*g*_uninfl_ = −0.64, ratio = 0.75) and ERBB4 (*g*_uninfl_ = −0.49, ratio = 0.99) were also constitutively suppressed.

Among inflammation-amplified genes, KYNU (*g*_infl_ = +2.22, *g*_uninfl_ = +0.45), GPX8 (+1.85, +0.70), and TGFBI (+1.72, +0.47) showed the largest differences in effect size between conditions.

At the pathway level, 122 of the 172 significant uninflamed pathways were also significant in the inflamed analysis, all with the same direction (Figure 4). The 86 shared upregulated pathways spanned the major immune and ECM clusters identified in the inflamed analysis, including interleukin-10 signaling (uninflamed NES = +2.07 vs inflamed +2.55), interferon *α*/*β* signaling (+1.69 vs +2.51), interleukin-4/13 signaling (+1.98 vs +2.43), complement cascade (+1.79 vs +2.22), PD-1 signaling (+1.98 vs +2.21), collagen formation (+1.47 vs +2.38), and integrin interactions (+1.91 vs +2.22). Neutrophil degranulation, on the other hand, was not significant in uninflamed tissue (NES = +1.28, *q* = 0.10). Among the 36 shared downregulated pathways, glucuronidation (uninflamed NES = −2.22 vs inflamed −2.20), ubiquinol biosynthesis (−1.91 vs −1.90), lipid metabolism (−1.56 vs −1.66), and coenzyme A biosynthesis (*−*1.48 vs −1.37) were all suppressed. Notably, the TCA cycle and respiratory electron transport (uninflamed NES = −0.52, *q* = 0.99), mitochondrial fatty acid *β*-oxidation (−0.93, *q* = 0.86), cristae formation (−0.53, *q* = 0.95), and formation of ATP by chemiosmotic coupling (−0.32, *q* = 0.99) were *not* significantly dysregulated in uninflamed tissue, indicating that these metabolic deficiencies are dependent on active inflammation. However, individual genes in some of these pathways (e.g., OPA1, IDH3A) were already significantly reduced, suggesting that the metabolism is partially compromised even in the absence of visible inflammation.

**Fig. 4.**
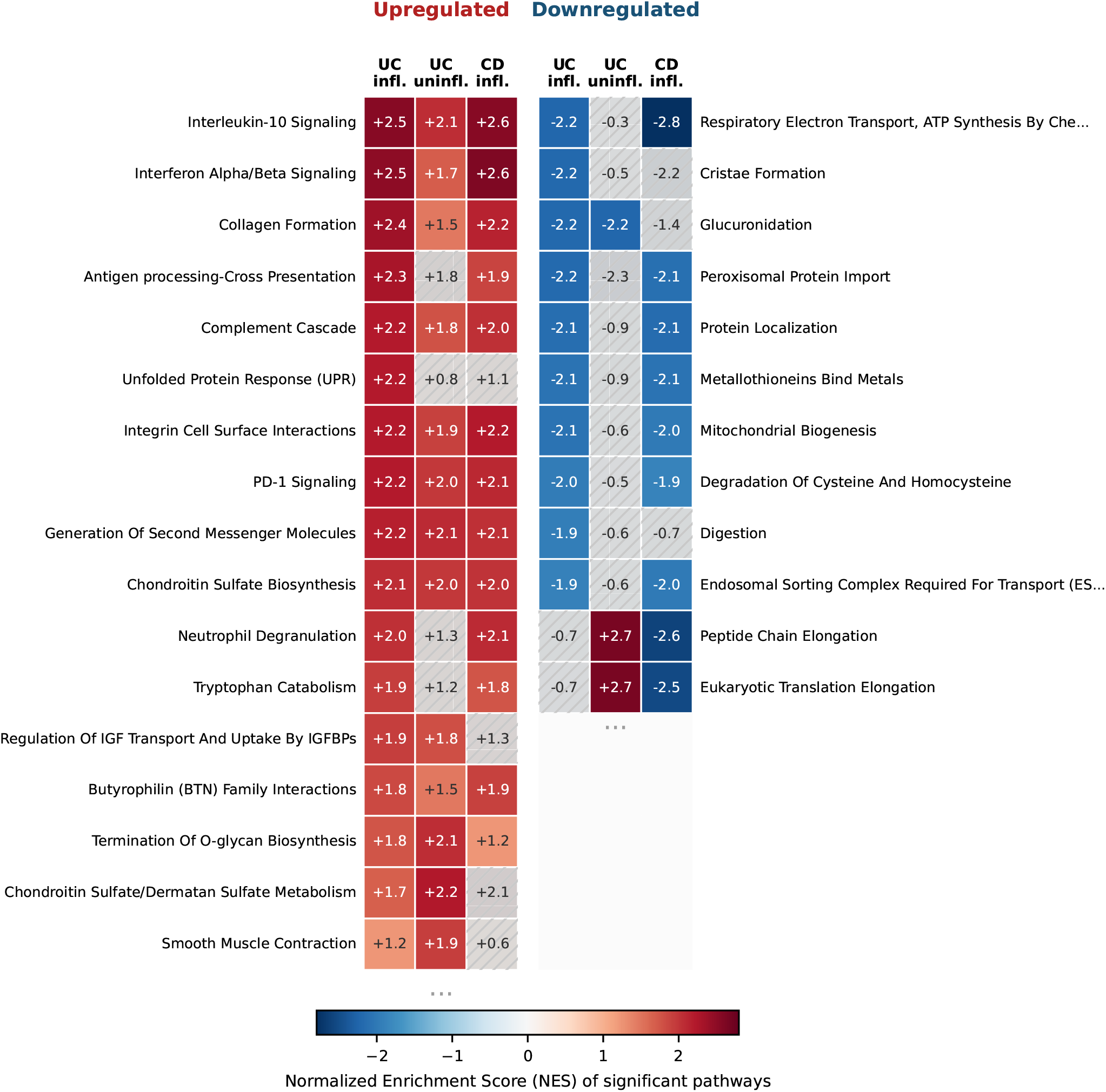
Three-way comparison of pathway enrichment across UC inflamed, UC uninflamed, and CD inflamed analyses. Each row is a cluster-representative pathway from the three analyses; colour encodes normalized enrichment score (red = upregulated in disease, blue = downregulated).

Among the 50 uninflamed-only pathways, ribosomal translation group — including eukaryotic translation elongation (NES = +2.66), peptide chain elongation (+2.65), SRP-dependent cotranslational targeting (+2.59), nonsense-mediated decay (+2.49) — was the overwhelmingly dominating signal. This translation upregulation was absent in inflamed UC, where eukaryotic translation elongation showed a weak, non-significant negative enrichment (NES = *−*0.69).

### Comparison with Crohn’s disease

8 datasets comparing inflamed CD to controls yielded 2,494 significantly dysregulated genes (1,245 up, 1,249 down), with a median |*g*| = 0.86 and *I*^2^ = 0.63. Gene-level effect sizes were highly correlated between UC and CD (Pearson *r* = 0.86 across 24,821 shared genes; Supplementary Tables S1 and S3), and 2,132 of the 2,494 CD-significant genes were also significant in UC, with all but one (CLCA1; down in UC, up in CD) showing the same direction of change.

Pathway-level results were similarly consistent (Figure 4). Of the 386 significantly enriched CD pathways, 288 (75%) were also significant in UC, all with the same direction of enrichment. Among these, 63 pathways were significant across all three disease-versus-control analyses (UC inflamed, UC uninflamed, and CD inflamed), dominated by ECM remodelling and collagen turnover, the complement cascade, and interferon/interleukin signaling. Further 225 pathways were shared between UC and CD but not significant in uninflamed UC, consistent with their classification as inflammation-dependent. They concentrated around immune/cytokine signaling, mitochondrial energy and lipid metabolism, growth factor/receptor signaling, cell death and autophagy, as well as oxidative stress.

The most prominent CD-specific finding was a group of 21 downregulated ribosomal and translation pathways among the 98 CD-only pathways, including Peptide Chain Elongation (NES = −2.59), Eukaryotic Translation Elongation (−2.52), and Selenocysteine Synthesis (−2.55). This group had no counterpart among the UC significant pathways, where Eukaryotic Translation Elongation was only weakly affected (NES = −0.69, *q* = 0.38). Interestingly, the same translation machinery was strongly *upregulated* in uninflamed UC (NES = +2.66), creating a three-way divergence: upregulated in uninflamed UC, neutral in inflamed UC, and strongly downregulated in CD.

### Direct comparison of UC and CD

The indirect comparison above used each disease’s deviation from controls to infer UC–CD differences. To validate these findings without relying on control samples, we performed a direct meta-analysis comparing inflamed UC biopsies with inflamed CD biopsies within the 6 datasets that contained both disease groups.

The direct comparison identified 1,813 significantly differentially expressed genes (*q* < 0.05; 845 higher in UC, 968 lower in UC relative to CD), with a median |*g*| = 0.55 and negligible heterogeneity (median *I*^2^ = 0.00). Gene-level effect sizes from the direct comparison were strongly correlated with the indirect difference (*g*_UC vs HC_ − *g*_CD vs HC_; Pearson *r* = 0.74 across 24,810 shared genes; Figure 5), confirming that the indirect approach reliably captures true UC–CD differences. At the pathway level, 493 pathways were significantly enriched, of which 85 passed the knee-point filter (44 higher in UC, 41 lower in UC). Louvain clustering grouped these into 22 communities. The downregulated (lower in UC than CD) pathway landscape was completely dominated by 4 clusters capturing the core metabolic signature: respiratory electron transport, TCA cycle, and cristae formation (18 pathways; mean NES = *−*2.14), mitochondrial fatty acid *β*-oxidation and peroxisomal metabolism (9 pathways; −1.98), peroxisomal lipid metabolism (6 pathways, −1.97), and glucuronidation (2 pathways; −2.10). These results confirm that the metabolic deficit identified in the disease-versus-control analyses is more severe in UC than in CD.

**Fig. 5.**
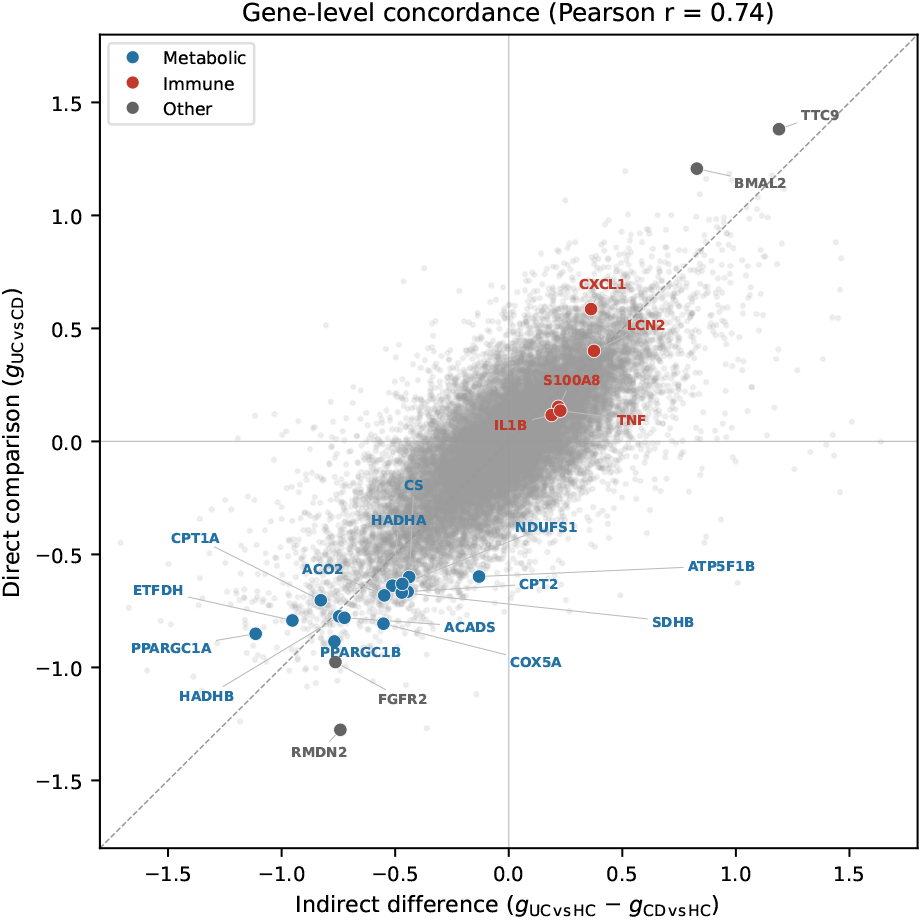
Concordance between indirect and direct UC–CD gene-level effect sizes. Each point represents one gene; the *x*-axis shows the indirect difference (*g*_UC vs HC_ *− g*_CD vs HC_) and the *y*-axis the direct comparison (*g*_UC vs CD_). High-lighted genes belong to metabolic (blue) or immune (red) categories. Pearson *r* = 0.74 across 24,810 shared genes.

Among the 12 upregulated clusters (higher in UC than CD), the largest one focused on translation, rRNA processing, and the unfolded protein response (16 pathways; mean NES = +1.95), including SRP-dependent cotranslational targeting (NES = +2.14), unfolded protein response (+2.13), and eukaryotic translation elongation (+1.92). This cluster directly validates the three-way translation divergence observed in the indirect analyses: translation pathways are higher in UC than in CD because CD strongly downregulates them while UC does not. A prefoldin- and tubulin-folding cluster was also elevated (5 pathways; NES = +2.11). Other notable upregulated clusters included collagen and ECM assembly pathways (9 pathways; NES = +1.97), complement regulation (3 pathways; NES = +1.94), and cell-cycle pathways including activation of ATR in response to replication stress (3 pathways; NES = +2.02).

Importantly, inflammation-shared pathways were not significant in the direct comparison: interferon *α*/*β* signaling (NES = +0.49, *q* = 0.69), PD-1 signaling (+0.68, *q* = 0.54), interleukin-10 signaling (+1.26, *q* = 0.16), and neutrophil degranulation (+0.87, *q* = 0.37) showed no UC–CD difference, consistent with both diseases activating similar immune responses.

At the gene level, the mitochondrial biogenesis regulators PPARGC1A (PGC-1*α*; *g* = −0.85, *q* = 1.0×10^*−*3^), PPARGC1B (PGC-1*β*; *g* = −0.89, *q* = 5.9×10^*−*5^), and ES-RRA (ERR*α*; *g* = −0.81, *q* = 2.5×10^*−*4^) were all significantly lower in UC than in CD. These transcription factors coordinate expression of genes involved in oxidative phosphorylation, fatty acid oxidation, and the TCA cycle.

## Discussion

This study integrated 14 independent microarray datasets from the Gene Expression Omnibus, spanning 9 array platforms and comprising 972 mucosal biopsy samples, to identify genes and pathways consistently dysregulated in UC across cohorts. To avoid often uncertain cross-study normalization, rather than pooling raw data across platforms, we computed effect sizes (Hedges’ *g*) and enrichment scores independently within each dataset and combined them through random-effects meta-analysis (REML). This design confines platform-specific biases within each study and allows heterogeneity to be quantified explicitly via the *I*^2^ statistic rather than masked by batch correction. At every analytical layer, we imposed conservative significance thresholds (*q* < 0.05, direction consistency ≥ 80%, presence in ≥ 50% of datasets) to ensure that reported results reflect real biological signal rather than artefacts of any single cohort. Crucially, each primary result was cross-validated by at least one complementary method: gene-level REML rankings were confirmed by non-parametric Robust Rank Aggregation (Spearman *ρ* = 0.89), pathway-level REML estimates were validated by Stouffer’s weighted *z*-method (91.3% concordance, 100% direction agreement), and UC–CD differences inferred indirectly through separate disease-versus-control analyses were validated by a direct within-dataset UC-versus-CD comparison (Pearson *r* = 0.74). To move from individual pathways to interpretable biological themes, we applied a data-driven knee-point filter to retain only the most strongly enriched pathways and then clustered them by shared leading-edge genes using the Louvain algorithm. The resulting pathway communities form the basis of the biological interpretation that follows. Four parallel analyses (inflamed UC versus controls, uninflamed UC versus controls, inflamed CD versus controls, and direct UC versus CD) were conducted through an identical pipeline, enabling systematic classification of each molecular change as constitutive or inflammation-dependent and as UC-specific or shared with CD. The most important results addressed in the discussion are presented in Table 5.

**Table 5.**
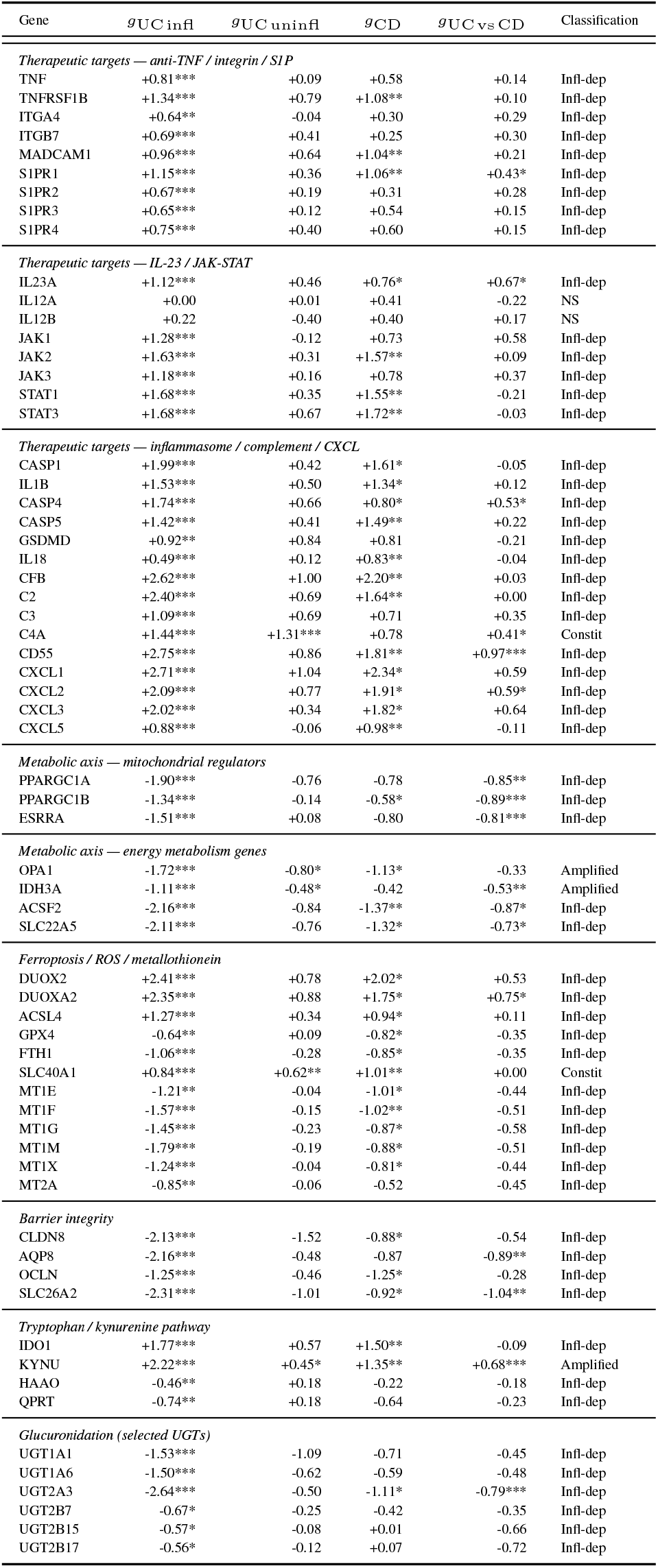
Cross-comparison gene-level effect sizes for key discussed genes. Each column shows the meta-analytic Hedges’ *g* from one of the four analyses; significance is indicated by stars (^*\**^*q* < 0.05, ^*\*\**^*q* < 0.01, ^*\*\*\**^*q* < 0.001). Classification: Constit = constitutive (significant in both, non-significant difference between effect sizes in inflamed and uninflamed, ratio 0.70–1.30); Amplified = inflammation-amplified (significant in both, significant difference between both); Infl-dep = inflammation-dependent (significant only in inflamed); NS = not significant in either. The UC vs CD column shows the direct comparison (positive = higher in UC).

### The inflammatory landscape

The upregulated transcriptomic landscape of inflamed UC mucosa is dominated by immune and inflammatory pathways, and the gene-level data align closely with the targets of every approved drug class. Tumour necrosis factor (TNF) and its receptor TN-FRSF1B were both significantly upregulated, yet TNF itself ranked outside the top 1,000 upregulated genes. This relatively weak transcriptional signal is consistent with the 20– 40% primary non-response rate to anti-TNF agents (12–14) and suggests that TNF-driven pathways are not the dominant factor in a substantial subset of patients. The integrin subunits ITGA4 and ITGB7, together with their endothelial ligand MAdCAM1, were consistently upregulated, confirming the enhanced lymphocyte trafficking addressed by vedolizumab (7).

The interleukin-23 axis provided a particularly informative pattern. IL23A (encoding the p19 subunit specific to IL-23) was significantly upregulated, whereas IL12A (p35, specific to IL-12; *g* = 0.00, *q* = 0.99) and IL12B (p40, shared between IL-12 and IL-23; *q* = 0.27) were not. This discrepancy may at least partly explain the growing body of evidence demonstrating the clinical superiority of selective IL-23 blockade over the concomitant inhibition of both IL-23 and IL-12 (10, 11). It may also explain the efficacy of anti-p19 antibodies (guselkumab, risankizumab, mirikizumab), even in patients who have failed ustekinumab therapy.

JAK-STAT signaling showed the strongest and most coherent immune signal, with JAK1–3, STAT1, and STAT3 all significantly upregulated. The breadth of activation across the entire cascade is consistent with the rapid therapeutic effect reported for tofacitinib, upadacitinib, and filgotinib (9), as these agents intercept the shared signaling hub rather than a single upstream cytokine. The S1P receptor family (S1PR1–4) was also upregulated, reflecting lymphocyte infiltration addressed by ozanimod and etrasimod (11), although with more moderate effect sizes.

Every therapeutic target examined was strictly inflammation-dependent, returning to baseline in uninflamed UC mucosa. This makes sense from a pharmacological point of view — immunosuppressants should target processes active during flares — but it also means that none of these targets address whatever renders the uninflamed mucosa vulnerable to the next flare. Moreover, while the immune signal dominates in terms of the number of upregulated genes, the downregulated landscape tells a different story: a coordinated collapse of mitochondrial energy metabolism.

### Metabolic disruption

The most striking non-immune finding was the simultaneous suppression of every layer of the mitochondrial energy production: cristae formation, fatty acid *β*-oxidation, the TCA cycle, and respiratory electron transport. These are not independent observations — they represent successive stages of the same energetic pipeline, from the structural scaffolding of the inner mitochondrial membrane through substrate oxidation to ATP generation. Their coordinated collapse can be explained by the fact that PPARGC1A (PGC-1*α*), PPARGC1B (PGC-1*β*), and ESRRA (ERR*α*) — the transcriptional co-activators that coordinate mitochondrial biogenesis, oxidative phosphorylation, and fatty acid oxidation (48) — were all significantly suppressed. This is consistent with established mechanisms linking inflammatory NF-*κ*B signaling to PGC-1*α* suppression (49), and our data confirm that the suppression is strictly inflammation-dependent — none of the 3 co-activators was significantly altered in uninflamed tissue, and all 4 downstream pathways were non-significant (all *q >* 0.85). Nonetheless, scattered metabolic genes were already reduced in uninflamed tissue, as detailed in the constitutive analysis below.

The energy deficit triggered a cascade of secondary failures. Metallothioneins were uniformly suppressed (strictly inflammation-dependent). Their loss removes both antioxidant and zinc supply, with direct consequences to barrier integrity, as intracellular zinc is required for the maintenance of tight junction proteins including occludin and claudin-3 (50). Simultaneously, DUOX2/DUOXA2 (responsible for generating ROS) was among the most strongly upregulated genes in the entire dataset (also inflammation-dependent). The collision of increased ROS production with decreased antioxidant capacity creates conditions favouring ferroptosis — a form of iron-dependent cell death driven by lipid peroxidation (51). Several findings support this: ACSL4, which channels polyunsaturated fatty acids into membrane phospholipids (52), was upregulated; GPX4 (the principal enzyme suppressing ferroptosis) and FTH1 (ferritin heavy chain which stores iron), were both downregulated. Moreover, SLC40A1 (ferroportin) was constitutively upregulated, indicating that iron handling is already altered before visible inflammation begins.

The consequences of these changes were severe: claudin-8, aquaporin-8, and occludin — core tight junction and transport components — were all significantly downregulated and inflammation-dependent. A breached barrier allows luminal antigens to access lamina propria, triggering further immune activation that exacerbates the metabolic deficiency, which accelerates barrier failure. This self-reinforcing cycle — energy deficit, oxidative stress, barrier breakdown, immune activation, deeper energy deficit — offers a mechanistic explanation for the clinical observation that UC flares escalate unless pharmacologically interrupted, and that remission achieved by immunosuppression alone is fragile as it breaks the cycle at the immune level but leaves the metabolic vulnerability unaddressed.

One pathway illustrates the potential direct connection between inflammation and metabolic collapse. IDO1, the rate-limiting enzyme of tryptophan catabolism induced by interferon-*γ* (53), was strongly upregulated, diverting tryptophan into the kynurenine pathway. KYNU, one of the top 20 upregulated genes, was the only enzyme in this pathway already significantly upregulated in uninflamed tissue (inflammation-amplified). However, the terminal enzymes HAAO and QPRT — required to complete the pathway to NAD+ (54) — were both downregulated. The result is a truncated pathway: inflammation drives tryptophan consumption, but the intermediates cannot be converted to NAD+, an essential cofactor for the TCA cycle. Tryptophan catabolism thus represents a potential molecular bridge through which immune activation directly deepens the metabolic deficit. Whether NAD+ depletion through this mechanism contributes independently to the energy deficit or is itself a downstream consequence of the broader metabolic collapse needs to be verified experimentally.

### Epithelial and stromal layers

An important interpretive caveat for the preceding sections is that bulk transcriptomics cannot assign signals to individual cell types. The metabolic signature — PPARGC1A suppression, UGT loss, barrier protein downregulation — involves genes predominantly expressed in colonocytes (55) (although may also reflect altered cell composition — fewer colonocytes and more infiltrating immune cells), whereas the collagen formation and ECM organisation cluster (15 pathways) involves genes predominantly expressed by stromal fibroblasts (56). These two layers are undergoing distinct but interacting processes within the same biopsy.

In the stroma, the transcriptomic picture suggests chaotic matrix turnover. Matrix metalloproteinases were upregulated — driven by neutrophils, macrophages, and activated fibroblasts — accelerating extracellular matrix degradation (57). Simultaneously, collagen biosynthesis, integrin and syndecan interactions, and chondroitin sulfate biosynthesis were all enriched, consistent with a fibroblast repair response attempting to compensate for the tissue damage (58). The net result is likely a disorganised matrix: fibrosis in some areas, erosion in others, and a basement membrane that cannot properly support the overlying epithelium. This reinforces barrier failure from below, complementing the tight junction loss from above described in the preceding section.

### Constitutive vulnerabilities and pre-inflammatory pathology

The analyses described so far have focused on inflamed UC mucosa where immune activation and metabolic collapse co-occur. The uninflamed UC analysis complements this picture by revealing a set of molecular alterations that are fully present in uninflamed tissue.

The most striking constitutive deficit was glucuronidation. At the pathway level, glucuronidation was equally suppressed in uninflamed and inflamed tissue (NES = −2.22 and −2.20, respectively) — a virtually identical signal. No individual UGT gene reached significance in the 5-dataset uninflamed analysis, but all 22 UGT genes tested had negative effect sizes, with 15 showing *g* < −0.3. GSEA detects this coordinated shift even when no single gene crosses the FDR threshold. The implication is that Phase II detoxification is already impaired in uninflamed UC mucosa: the tissue enters each inflammatory episode unable to efficiently conjugate xenobiotics and reactive metabolites, including bilirubin — an endogenous antioxidant processed by UGT1A1 (59). Importantly, this deficiency is not downstream of the mitochondrial energy collapse described above, since the energy collapse is inflammation-dependent and glucuronidation is already suppressed before inflammation is present. Beyond detoxification, ubiquinol and coenzyme A biosynthesis, as well as general lipid metabolism were constitutively suppressed at the pathway level, and individual mitochondrial genes (e.g., OPA1, IDH3A) were already reduced, indicating metabolic vulnerability before visible inflammation.

Translation pathways revealed an even more unexpected pattern. The three-way divergence described in the Results — strongly upregulated in uninflamed UC, neutral in inflamed UC, strongly downregulated in CD — suggests that uninflamed UC mucosa already has some underlying, unidentified stress to which the cells try to respond. Inflammation overwhelms this response — likely because the energy deficit removes the ATP needed to sustain ribosomal activity — and CD suppresses it through what may be a different mechanism. The presence of this repair response in tissue classified as uninflamed implies that the primary stress in UC precedes the immune response.

Other constitutive changes reinforce this interpretation. C4A — a complement component — was the only constitutive complement gene, while all other complement components (CFB, C2, C3, CD55) were strictly inflammation-dependent. The innate immune system appears to be partially active before visible inflammation takes place. SLC40A1 (ferroportin) was constitutively upregulated, which indicates altered iron homeostasis in uninflamed mucosa that requires compensatory iron export. This could suggest a vulnerability to ferroptosis which then gets exacerbated by active inflammation through suppressing GPX4 and FTH1. KYNU, the kynurenine pathway enzyme, had a small but significant uninflamed effect (inflammation-amplified), suggesting that the tryptophan diversion described earlier is weakly active even at baseline.

Taken together, these findings suggest a two-layer model of UC pathology. The first layer is constitutive: impaired detoxification, weakened oxidative defences, partially heightened complement, altered iron handling, translation machinery activated in a repair response, and scattered metabolic genes already trending downward. The second layer is inflammation-dependent: full PPARGC1A/B/ESRRA suppression triggers mitochondrial energy collapse and metallothioneins loss, DUOX2/DUOXA2 generates reactive oxygen species, barrier proteins are degraded, tryptophan is diverted, and the vicious cycle locks in. It seems that the inflammation does not create the vulnerability from scratch — it exploits a pre-existing one. This distinction has direct therapeutic implications, discussed below.

Although the constitutive layer observations about UC aetiology are speculative, they narrow the field of possibilities. The primary trigger appears to be upstream of both inflammation and the full metabolic collapse. Virtually all immune pathways are inflammation-dependent, making the immune system an unlikely primary driver. Our analysis suggests that the constitutive vulnerabilities might be metabolic and detoxification-related, pointing toward environmental or dietary factors.

### Differentiation from Crohn’s disease

The comparison with CD enhances the specificity of the findings above. At the immune level, the two diseases are remarkably similar. Gene-level effect sizes are strongly correlated (*r* = 0.86) and the major immune pathways were present in both diseases, with almost none reaching significance in the direct UC-versus-CD comparison (except for IL-2 and IL-27 which were higher in UC), consistent with both diseases activating a shared inflammatory response. This convergence explains why the same immunosuppressive agents (anti-TNF, JAK inhibitors) are effective in both conditions.

Metabolism is the most characteristic feature distinguishing UC from CD. Although significantly disrupted in both diseases, respiratory electron transport, fatty acid *β*-oxidation, and glucuronidation were all significantly more suppressed in UC, and the three master regulators PPARGC1A, PPARGC1B, and ESRRA were also lower. This higher severity likely reflects the unique metabolic reliance of the colonic epithelium and the anatomical distribution of the diseases. Colonocytes derive up to 70% of their energy from the mitochondrial *β*-oxidation of microbiome-derived short-chain fatty acids, primarily butyrate (60). Because UC is continuous and strictly confined to the colonic mucosa, the inflammatory damage uniformly destroys the epithelial layer where this metabolic profile is present. Furthermore, the loss of butyrate transporters and butyrate-producing bacteria in UC starves the colonocytes of their primary source of energy, forcing a more profound shutdown of the *β*-oxidation and respiratory pathways than in the patchy, transmural inflammation characteristic of CD.

Translation pathways were significantly higher in UC than in CD in the direct comparison, consistent with the three-way pattern identified in the indirect analyses. This can be explained under the two-layer model: UC mucosa has a constitutive translational repair response that CD mucosa does not, possibly because the underlying stress (or the response to it) differs between the two diseases. The loss of this response in inflamed UC (where it is not dysregulated) and its active suppression in CD may reflect different downstream consequences of the shared inflammatory response acting on epithelia with distinct starting points.

### Therapeutic implications

The findings of this study suggest two categories of unmet therapeutic need.

The first is a set of upregulated immune pathways for which no approved UC therapy currently exists. The inflamma-some axis — CASP1, IL1B, CASP4, CASP5, GSDMD, IL18 — was consistently activated across cohorts. This activation was completely inflammation-dependent, representing a coordinated pyroptotic signaling system that is not targeted by any approved biologic or small molecule for UC. The complement system (CFB, C2, C3, and C4A) was similarly prominent, with C4A notable as one of the few immune genes already upregulated before inflammation, suggesting that complement priming may contribute to flare susceptibility. CXCL chemokines (CXCL1–3, CXCL5) — key neutrophil recruitment signals — were among the most strongly upregulated genes and are likewise not directly targeted by current therapies. Whether these pathways represent viable drug targets requires clinical investigation, but the transcriptomic evidence for their consistent activation across independent cohorts is strong.

The second category — and arguably the more consequential — is the metabolic dimension, which is entirely unaddressed by current therapies. No approved UC drug targets the mitochondrial energy deficiency, the glucuronidation collapse, or the oxidative stress described above. The two-layer model suggests several potential intervention strategies. PPAR*γ* agonists promote mitochondrial fatty acid *β*-oxidation in colonocytes and could in principle counteract the inflammation-driven suppression of mitochondrial energy production (61). NAD+ precursors (such as nicotinamide riboside) could bypass the truncated kynurenine pathway and restore cofactor availability for the TCA cycle. Butyrate or short-chain fatty acid supplementation would directly address the colon’s preferred fuel deficiency. Mitochondrial-targeted antioxidants could mitigate the ferroptotic vulnerability created by the convergence of DUOX2 activation, GPX4 suppression, and metallothionein loss. These are speculative suggestions that require experimental validation, but they address mechanisms that are well supported by the transcriptomic data.

Perhaps the most consequential implication of the two-layer model is the concept of a therapeutic window between flares. The constitutive vulnerabilities — impaired detoxification, weakened antioxidant reserves, altered iron handling — persist during clinical remission. This means the mucosa remains metabolically fragile even when inflammation has been pharmacologically suppressed, which may explain the high relapse rates observed even under maintenance therapy. One immediate clinical implication concerns iron supplementation for anemia. Current guidelines recommend intravenous iron as first-line treatment in active UC but permit oral iron during remission (62). However, unabsorbed luminal iron can promote ferroptosis in epithelial cells (63). Combined with the constitutive upregulation of SLC40A1 (ferroportin) identified in this analysis, these findings raise the possibility that intravenous iron may be preferable to oral iron even outside active disease. This possibility should be evaluated in prospective clinical studies.

Current UC therapies focus on suppressing the inflammatory response, but do not address the epithelial substrate on which inflammation acts. If the constitutive layer could be addressed during remission — by restoring glucuronidation and antioxidant capacity or supporting mitochondrial function — it might be possible to raise the threshold at which inflammation triggers the vicious cycle of metabolic collapse.

### Limitations

Several limitations should be considered when interpreting these findings. First, all datasets were generated on microarray platforms. Microarrays have limited dynamic range and probe-dependent gene coverage, meaning that genes with low expression and transcripts absent from platform annotations may be missed. Second, bulk tissue transcriptomics conflates signal from epithelial cells, stromal fibroblasts, and infiltrating immune cells. As discussed above, some apparent downregulation of colonocyte-expressed genes may reflect altered cell composition rather than true transcriptional reprogramming, and single-cell resolution would be needed to definitively assign signals to specific cell types. Third, the constitutive analysis relied on only 5 datasets with uninflamed UC samples, yielding substantially reduced statistical power (165 significant genes versus 7,727 in the inflamed analysis). Gene-level effects that did not reach significance — such as individual UGT genes or CLDN8 — may represent real biological changes that the current sample size cannot detect. Moreover, we cannot control for medication history or exclude subclinical inflammation in samples labelled as uninflamed, both of which could confound the constitutive signal. Fourth, several of the most strongly dysregulated genes — including BACE2, DPP10, and PDE6A — fall outside annotated Reactome pathways and were therefore not captured by the pathway-level analysis. Finally, and most importantly, the study is observational. The proposed two-layer model and the vicious cycle connecting energy deficit, oxidative stress, and barrier failure are consistent with the transcriptomic data but remain hypotheses. Establishing causality — particularly whether the constitutive vulnerabilities precede and predispose to inflammation, or are themselves consequences of prior subclinical disease — will require additional experimentation and interventional studies.

