## Supplementary material for "Large-scale transcriptomic meta-analysis identifies novel therapeutic targets for ulcerative colitis"

Piernik et al.

#### Contents

1 Supplementary Figures 2

2 Supplementary Tables 5

### 1 Supplementary Figures

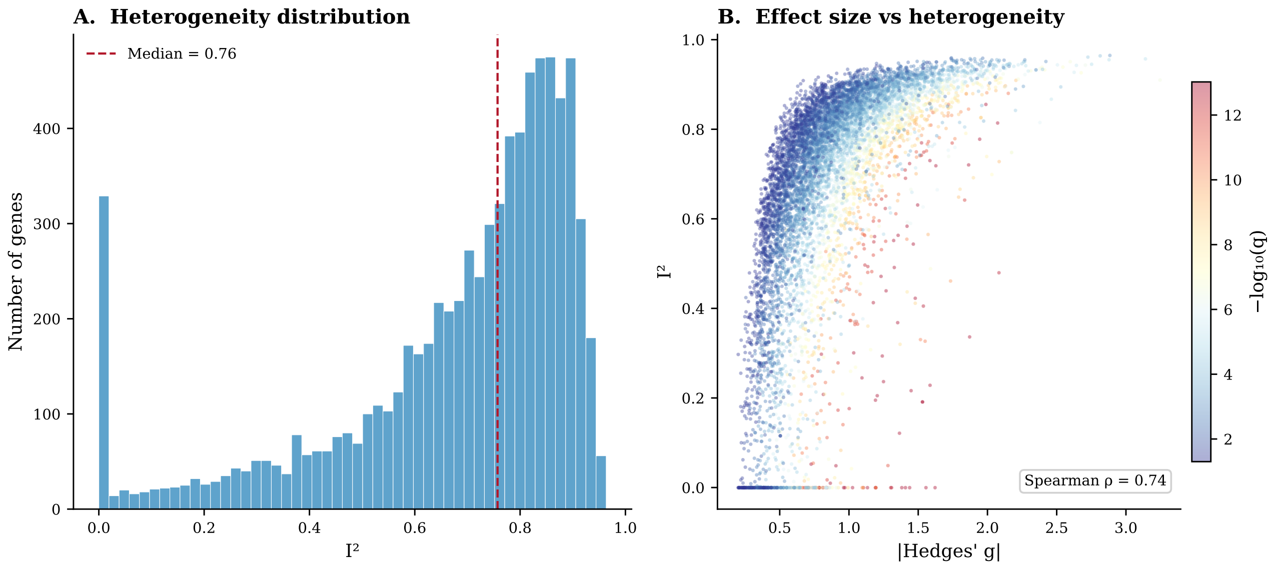


Figure S1: **Heterogeneity diagnostics for the inflamed UC vs control gene-level meta-analysis.** **(A)** Distribution of $I^2$ across 7,727 significantly dysregulated genes. The median $I^2$ of 0.76 reflects variation in effect-size magnitude across the 12 datasets and 9 microarray platforms, rather than inconsistency in direction (controlled by the $\geq 80\%$ direction consistency filter). **(B)** Relationship between absolute effect size ($|g|$) and heterogeneity ($I^2$). Colour encodes statistical significance ($-\log_{10} q$). Genes with large effects tend to have high $I^2$ (Spearman $\rho = 0.74$), while genes ranking highest by $q$-value have low $I^2$, reflecting complementary properties of the two ranking criteria.

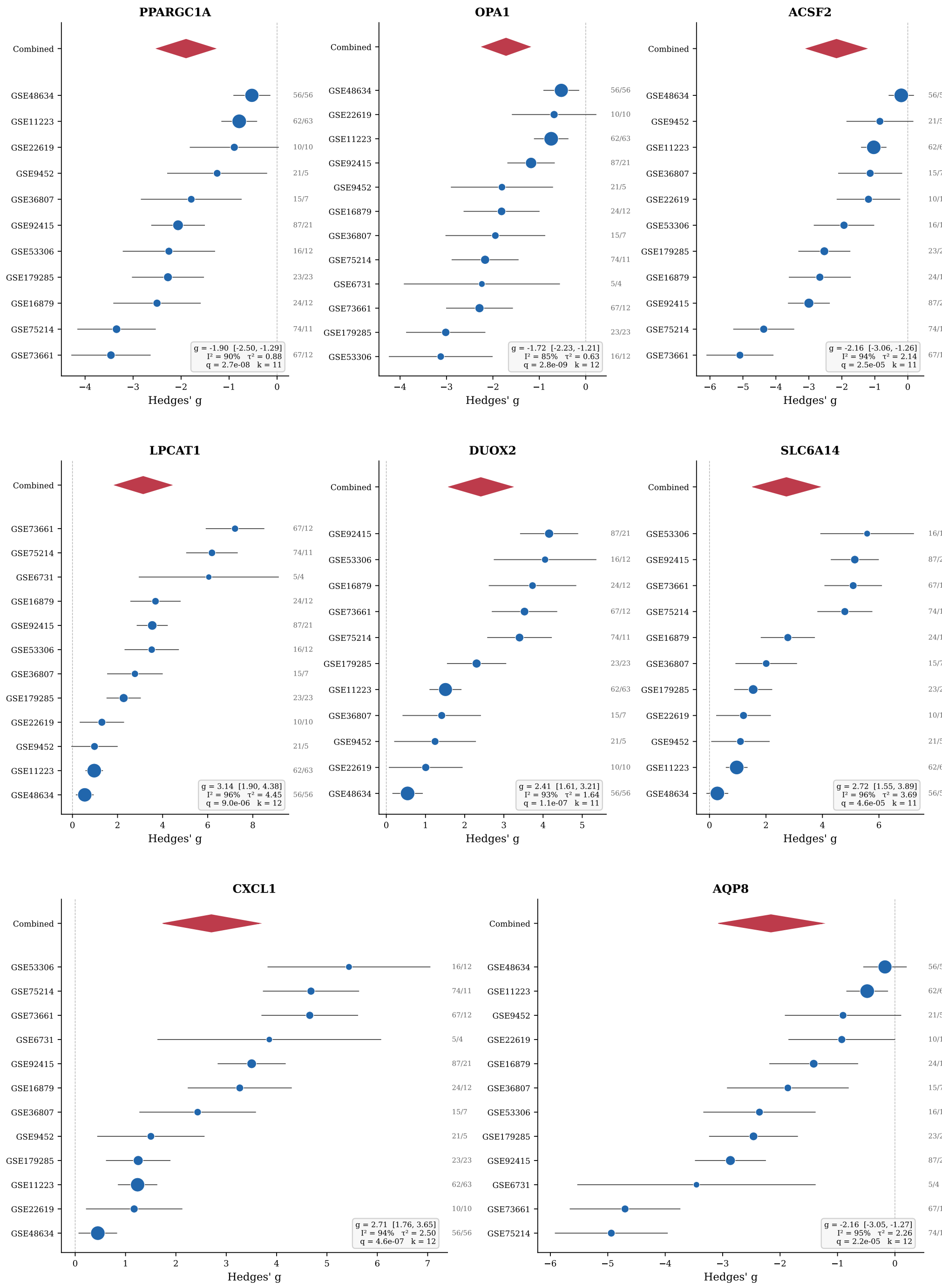


Figure S2: **Forest plots for representative genes from the inflamed UC vs control meta-analysis.** Each panel shows per-dataset effect sizes (Hedges' $g$, blue circles scaled by inverse variance) with 95% confidence intervals, and the random-effects combined estimate (red diamond). Numbers at right indicate sample sizes ($n_{\mathrm{UC}}/n_{\mathrm{Ctrl}}$). **Top row:** PPARGC1A, OPA1, and ACSF2 (downregulated metabolic genes). **Middle row:** LPCAT1, DUOX2, and SLC6A14 (top upregulated genes). **Bottom row:** CXCL1 (top upregulated immune) and AQP8 (downregulated barrier gene).

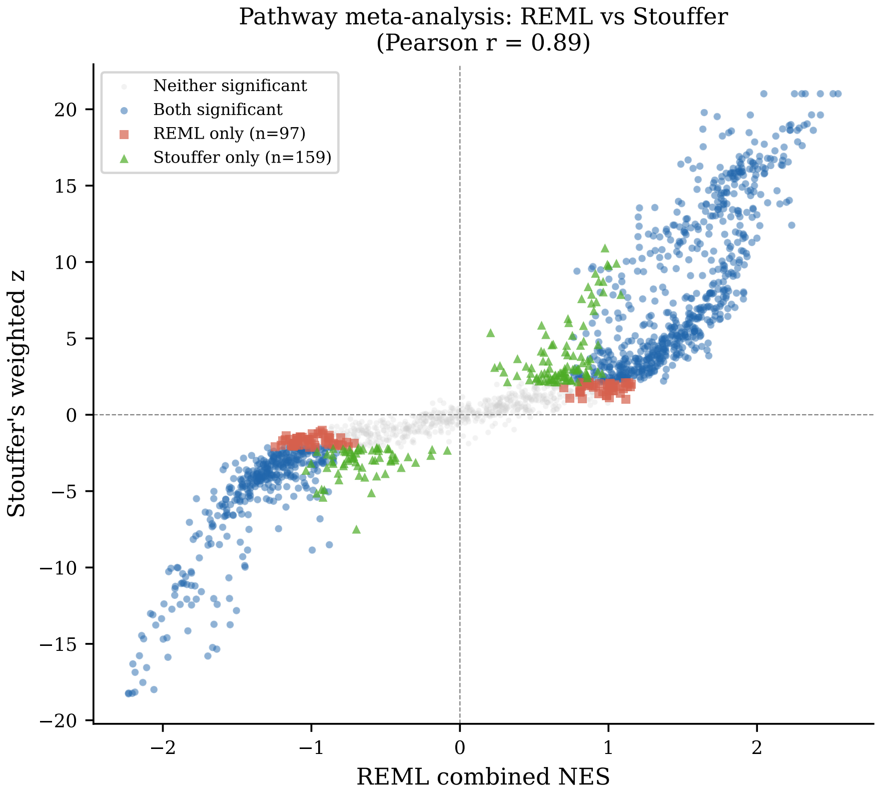


Figure S3: **Sensitivity analysis: REML random-effects vs Stouffer's weighted $z$-method for pathway-level meta-analysis.** Each point represents one Reactome pathway. The $x$-axis shows the REML combined normalized enrichment score (NES); the $y$-axis shows Stouffer's weighted $z$-statistic (weights $= \sqrt{n}$). Blue points are significant by both methods ($q < 0.05$); red squares are REML-only; green triangles are Stouffer-only; grey points are non-significant by both. 91.3% of REML-significant pathways were confirmed by the Stouffer method, with 100% direction concordance and Pearson $r = 0.89$, supporting the validity of the NES variance approximation used in the REML framework.

#### 2 Supplementary Tables

##### Tables S1–S4: Gene-level meta-analysis results

Full gene-level meta-analysis results for all four comparisons are provided as separate CSV files:

- **Table S1**: UC inflamed vs healthy controls (7,727 significant genes)
- **Table S2**: UC uninflamed vs healthy controls (165 significant genes)
- **Table S3**: CD inflamed vs healthy controls (2,494 significant genes)
- **Table S4**: UC vs CD direct comparison (1,831 significant genes)

Each file contains: gene symbol, number of datasets, Hedges' $g$, standard error, 95% confidence interval, $p$-value, $q$-value (Benjamini–Hochberg), $\tau^2$, $I^2$, direction of effect, and direction consistency ratio.

##### Tables S5–S8: Pathway-level meta-analysis results

Significant pathway-level meta-analysis results ($q < 0.05$) for all four comparisons are provided as separate CSV files:

- **Table S5**: UC inflamed vs healthy controls (828 significant pathways)
- **Table S6**: UC uninflamed vs healthy controls (172 significant pathways)
- **Table S7**: CD inflamed vs healthy controls (386 significant pathways)
- **Table S8**: UC vs CD direct comparison (493 significant pathways)

Each file contains: Reactome pathway term, number of datasets, combined NES, standard error, 95% CI, $p$-value, $q$-value, $\tau^2$, $I^2$, direction, direction ratio, and core leading-edge genes.

#### Table S9: Pathway clusters — UC inflamed vs healthy controls

| Cluster representative | Pathways | Mean NES |
|---|---|---|
| Unfolded Protein Response (UPR) | 3 | +2.19 |
| Complement Cascade | 3 | +2.17 |
| Collagen Formation | 15 | +2.14 |
| Interleukin-10 Signaling | 26 | +2.12 |
| PD-1 Signaling | 5 | +2.11 |
| Antigen processing-Cross Presentation | 20 | +2.07 |
| Chondroitin Sulfate Biosynthesis | 1 | +2.07 |
| Neutrophil Degranulation | 4 | +1.96 |
| Tryptophan Catabolism | 1 | +1.93 |
| Regulation Of IGF Transport And Uptake By IGFBPs | 1 | +1.93 |
| Glucuronidation | 2 | −2.14 |
| Cristae Formation | 11 | −2.10 |
| Metallothioneins Bind Metals | 2 | −2.06 |
| Peroxisomal Protein Import | 8 | −2.05 |
| Digestion | 1 | −1.94 |
| Degradation Of Cysteine And Homocysteine | 2 | −1.94 |

Table S9: Pathway clusters from the inflamed UC vs healthy controls analysis. Each row is a community of co-enriched Reactome pathways identified by Louvain clustering on shared leading-edge genes. Positive NES = upregulated in inflamed UC relative to controls; negative NES = downregulated.

#### Table S10: Pathway clusters — uninflamed UC vs healthy controls

| Cluster representative | Pathways | Mean NES |
|---|---|---|
| Eukaryotic Translation Elongation | 14 | +2.44 |
| Chondroitin Sulfate/Dermatan Sulfate Metabolism | 2 | +2.10 |
| Interleukin-10 Signaling | 2 | +2.02 |
| Generation Of Second Messenger Molecules | 5 | +2.02 |
| Termination Of O-glycan Biosynthesis | 2 | +2.01 |
| Smooth Muscle Contraction | 1 | +1.92 |
| Integrin Cell Surface Interactions | 1 | +1.91 |
| Glucuronidation | 1 | −2.22 |

Table S10: Pathway clusters from the uninflamed UC vs healthy controls analysis. Each row is a community of co-enriched Reactome pathways identified by Louvain clustering on shared leading-edge genes. Positive NES = upregulated in uninflamed UC relative to controls; negative NES = downregulated.

#### Table S11: Pathway clusters — CD inflamed vs healthy controls

| Cluster representative | Pathways | Mean NES |
|---|---|---|
| Interferon Alpha/Beta Signaling | 17 | +2.10 |
| Interleukin-10 Signaling | 14 | +2.09 |
| Integrin Cell Surface Interactions | 11 | +2.06 |
| Chondroitin Sulfate Biosynthesis | 1 | +2.03 |
| Butyrophilin (BTN) Family Interactions | 1 | +1.90 |
| Respiratory Electron Transport, ATP Synthesis By Chemiosmotic Coupling, Heat Production By Uncoupling Proteins | 6 | −2.46 |
| Peptide Chain Elongation | 16 | −2.39 |
| Protein Localization | 2 | −2.13 |
| Metallothioneins Bind Metals | 2 | −2.05 |
| Mitochondrial Biogenesis | 1 | −1.98 |
| Endosomal Sorting Complex Required For Transport (ESCRT) | 1 | −1.98 |

Table S11: Pathway clusters from the CD inflamed vs healthy controls analysis. Each row is a community of co-enriched Reactome pathways identified by Louvain clustering on shared leading-edge genes. Positive NES = upregulated in CD relative to controls; negative NES = downregulated.

#### Table S12: Pathway clusters — UC vs CD direct comparison

| Cluster representative | Pathways | Mean NES |
|---|---|---|
| Prefoldin Mediated Transfer Of Substrate To CCT/TriC | 5 | +2.11 |
| Activation Of ATR In Response To Replication Stress | 3 | +2.02 |
| POLB-Dependent Long Patch Base Excision Repair | 1 | +2.00 |
| Assembly Of Collagen Fibrils And Other Multimeric Structures | 9 | +1.97 |
| SRP-dependent Cotranslational Protein Targeting To Membrane | 16 | +1.95 |
| Regulation Of Complement Cascade | 3 | +1.94 |
| Early Phase Of HIV Life Cycle | 1 | +1.92 |
| RAF-independent MAPK1/3 Activation | 1 | +1.90 |
| tRNA Modification In Nucleus And Cytosol | 1 | +1.86 |
| Nuclear Events (Kinase And Transcription Factor Activation) | 1 | +1.85 |
| Defective Factor VIII Causes Hemophilia A | 1 | +1.82 |
| STING Mediated Induction Of Host Immune Responses | 2 | +1.80 |
| Respiratory Electron Transport | 18 | −2.14 |
| Glucuronidation | 2 | −2.10 |
| Endosomal/Vacuolar Pathway | 1 | −1.98 |
| Mitochondrial Fatty Acid Beta-Oxidation | 9 | −1.98 |
| Peroxisomal Lipid Metabolism | 6 | −1.97 |
| Degradation Of Cysteine And Homocysteine | 1 | −1.93 |
| Mitochondrial Calcium Ion Transport | 1 | −1.88 |
| Digestion | 1 | −1.88 |
| Budding And Maturation Of HIV Virion | 1 | −1.87 |
| Branched-chain Amino Acid Catabolism | 1 | −1.86 |

Table S12: Pathway clusters from the UC vs CD direct comparison analysis. Each row is a community of co-enriched Reactome pathways identified by Louvain clustering on shared leading-edge genes. Positive NES = upregulated in UC relative to CD; negative NES = downregulated in UC relative to CD.

#### Table S13: Constitutive genes

Genes significant in both the inflamed and uninflamed UC vs control meta-analyses, with no significant difference between conditions (Wald $q_\Delta \geq 0.05$) and effect-size ratio within 0.70–1.30.

Table S13: Constitutive genes (27 genes: 17 upregulated, 10 downregulated). Ratio = $g_{\text{uninfl}}/g_{\text{infl}}$.

| Gene | $g_{\text{infl}}$ | $g_{\text{uninfl}}$ | Ratio | $\Delta g$ | $q_\Delta$ |
|---|---|---|---|---|---|
| C4A | +1.44 | +1.31 | 0.91 | +0.12 | 0.916 |
| SHANK3 | +1.06 | +1.05 | 0.98 | +0.02 | 0.985 |
| EMB | +0.82 | +1.01 | 1.23 | −0.19 | 0.874 |
| SHC2 | +1.08 | +0.98 | 0.91 | +0.10 | 0.931 |
| NR2F1 | +0.96 | +0.88 | 0.92 | +0.08 | 0.932 |
| KLK12 | +0.94 | +0.77 | 0.81 | +0.17 | 0.819 |
| AMPD3 | +0.98 | +0.72 | 0.74 | +0.26 | 0.556 |
| IRX2 | +0.63 | +0.71 | 1.13 | −0.08 | 0.906 |
| SPARCL1 | +0.99 | +0.70 | 0.70 | +0.29 | 0.600 |
| SLC49A4 | +0.66 | +0.68 | 1.03 | −0.02 | 0.976 |
| SLC40A1 | +0.84 | +0.62 | 0.74 | +0.22 | 0.647 |
| FILIP1 | +0.76 | +0.61 | 0.80 | +0.15 | 0.836 |
| C1QTNF5 | +0.73 | +0.53 | 0.73 | +0.20 | 0.704 |
| KCNS3 | +0.57 | +0.50 | 0.88 | +0.07 | 0.914 |
| C12orf75 | +0.37 | +0.48 | 1.30 | −0.11 | 0.813 |
| FAM229B | +0.60 | +0.47 | 0.78 | +0.13 | 0.811 |
| CCL21 | +0.62 | +0.46 | 0.74 | +0.16 | 0.750 |

| Gene | $g_{\text{infl}}$ | $g_{\text{uninfl}}$ | Ratio | $\Delta g$ | $q_\Delta$ |
|---|---|---|---|---|---|
| MAP3K13 | −0.87 | −0.77 | 0.89 | −0.10 | 0.918 |
| PHLDB3 | −0.74 | −0.70 | 0.95 | −0.04 | 0.971 |
| WDR1 | −0.57 | −0.69 | 1.21 | +0.12 | 0.875 |
| DLGAP1 | −0.55 | −0.66 | 1.20 | +0.11 | 0.869 |
| PANK4 | −0.82 | −0.65 | 0.80 | −0.17 | 0.812 |
| GLIPR2 | −0.85 | −0.64 | 0.75 | −0.21 | 0.733 |
| ARHGEF26 | −0.67 | −0.51 | 0.76 | −0.16 | 0.721 |
| ERBB4 | −0.49 | −0.49 | 0.99 | −0.00 | 0.997 |
| ZNF334 | −0.55 | −0.49 | 0.88 | −0.07 | 0.920 |
| ELMOD1 | −0.51 | −0.48 | 0.94 | −0.03 | 0.961 |

#### Table S14: Inflammation-amplified genes

Genes significant in both the inflamed and uninflamed UC vs control meta-analyses whose effect sizes differed significantly between conditions (Wald $q_{\Delta} < 0.05$).

Table S14: Inflammation-amplified genes (21 genes). $\Delta g = g_{\text{infl}} - g_{\text{uninfl}}$.

| Gene | $g_{\text{infl}}$ | $g_{\text{uninfl}}$ | Ratio | $\Delta g$ | $q_{\Delta}$ |
|---|---|---|---|---|---|
| KYNU | +2.22 | +0.45 | 0.20 | +1.77 | 0.002 |
| TGFBI | +1.72 | +0.47 | 0.28 | +1.25 | 0.001 |
| ARHGEF9 | −1.65 | −0.48 | 0.29 | −1.17 | 0.012 |
| GPX8 | +1.85 | +0.70 | 0.38 | +1.15 | 0.012 |
| PDPN | +1.59 | +0.46 | 0.29 | +1.14 | 0.002 |
| GLTP | −1.62 | −0.53 | 0.33 | −1.09 | 0.019 |
| GXYLT2 | −1.84 | −0.76 | 0.41 | −1.08 | 0.047 |
| BLVRA | +1.53 | +0.46 | 0.30 | +1.07 | 0.007 |
| PTAFR | +1.52 | +0.49 | 0.32 | +1.03 | 0.002 |
| CHKA | −1.50 | −0.48 | 0.32 | −1.02 | 0.028 |
| SLC6A6 | +1.45 | +0.44 | 0.30 | +1.01 | 0.022 |
| ITGB2 | +1.37 | +0.45 | 0.33 | +0.92 | 0.009 |
| CTSK | +1.39 | +0.50 | 0.36 | +0.89 | 0.027 |
| PSAT1 | +1.36 | +0.51 | 0.38 | +0.85 | 0.020 |
| RNF183 | +1.24 | +0.49 | 0.40 | +0.74 | 0.015 |
| CHRDL2 | +1.22 | +0.47 | 0.39 | +0.74 | 0.036 |
| AGT | +1.33 | +0.63 | 0.47 | +0.70 | 0.049 |
| VNN2 | +1.16 | +0.46 | 0.40 | +0.69 | 0.040 |
| BCL6 | +1.20 | +0.50 | 0.42 | +0.69 | 0.021 |
| PTGDS | +1.18 | +0.51 | 0.44 | +0.66 | 0.026 |
| IDH3A | −1.11 | −0.48 | 0.43 | −0.64 | 0.028 |
